# Hepatocyte-specific loss of LAP2α protects against diet-induced hepatic steatosis and steatohepatitis in male mice

**DOI:** 10.1101/2022.03.18.484917

**Authors:** Kapil K. Upadhyay, Eun-Young K. Choi, Roland Foisner, M. Bishr Omary, Graham F. Brady

**Affiliations:** Division of Gastroenterology and Hepatology, Department of Internal Medicine, University of Michigan, Ann Arbor, Michigan; Department of Pathology, University of Michigan, Ann Arbor, Michigan; Max Perutz Labs, Medical University of Vienna, Vienna Biocenter Campus (VBC), Vienna, Austria; Robert Wood Johnson Medical School and the Center for Advanced Biotechnology and Medicine, Rutgers University, Piscataway, New Jersey

**Keywords:** LAP2α, lamin A/C, hepatic steatosis, steatohepatitis, C57BL/6J mice, high fat diet

## Abstract

There is increasing evidence for the importance of the nuclear envelope in lipid metabolism, nonalcoholic fatty liver disease (NAFLD), and nonalcoholic steatohepatitis (NASH). Human mutations in *LMNA*, encoding A-type nuclear lamins, cause early-onset insulin resistance and NASH, while hepatocyte-specific deletion of *Lmna* predisposes to NASH with fibrosis in male mice. Given that variants in the gene encoding LAP2α, a nuclear protein that regulates lamin A/C, were previously identified in patients with NAFLD, we sought to determine the role of LAP2α in NAFLD using a mouse genetic model. Hepatocyte-specific *Lap2a-knockout* (*Lap2α*^(ΔHep)^) mice and littermate controls were fed normal chow or high-fat diet (HFD) for 8 weeks or 6 months. In contrast to what was observed with hepatocyte-specific *Lmna* deletion, male *Lap2α*^(ΔHep)^ mice showed no increase in hepatic steatosis or NASH compared to controls. Rather, *Lap2α*^(ΔHep)^ mice demonstrated reduced hepatic steatosis, particularly after long-term HFD, with decreased susceptibility to diet-induced NASH. Accordingly, whereas pro-steatotic genes *Cidea, Mogat1*, and *Cd36* were upregulated in *Lmnα*-KO mice, they were downregulated in *Lap2α*^(ΔHep)^ mice, and there was a trend toward decreases in pro-inflammatory and pro-fibrotic genes. These data indicate that hepatocyte-specific *Lap2α* deletion protects against hepatic steatosis and NASH in mice; therefore, LAP2α might represent a potential therapeutic target in human NASH.

**Brief Summary:** Loss of LAP2α in mouse hepatocytes protected against diet-induced hepatic steatosis and NASH.

## Introduction

Nonalcoholic fatty liver disease (NAFLD) is a clinical condition defined by excess fat (more than 5% by weight or volume) deposition in the liver without excessive alcohol consumption or steatogenic medication use (1). The spectrum of NAFLD ranges from simple steatosis to nonalcoholic steatohepatitis (NASH), which can lead to progressive fibrosis, cirrhosis, and hepatocellular carcinoma (2–5). The rising global prevalence of NAFLD, in parallel with obesity, may be partly explained by diet and sedentary lifestyle (6). However, environmental factors and genetic risk are also vital contributors to the observed variations in NAFLD occurrence among populations (7–9). Importantly, effective medical treatment for NAFLD/NASH is currently an unmet need, partly due to an incomplete understanding of its pathogenesis (10).

Laminopathies are a group of rare diseases caused by mutations in genes encoding proteins of the nuclear lamina and nuclear envelope (11–13). The nuclear lamina is a dense multi-protein network inside the nucleus at the inner nuclear membrane surface and is primarily composed of nuclear intermediate filament proteins known as A- and B-type lamins, as well as their associated proteins (14, 15). It is a structural and functional link between the cytoskeleton and heterochromatin and regulates DNA replication, transcription, cell cycle, cellular differentiation, and apoptosis (16–19). Notably, laminopathies include lipodystrophy syndromes that are characterized by insulin resistance, hypertriglyceridemia, and hepatic steatosis, typically with NASH and progressive fibrosis (13, 20, 21).

Recent animal and human studies have supported a direct causal and hepatocyte-autonomous role for nuclear envelope-related mutations in diseases of lipid homeostasis and metabolism. For example, mutation of the gene encoding lamin B receptor (LBR), an inner nuclear membrane protein, is associated with impaired cholesterol synthesis (22). Additionally, hepatocyte-specific lamina-associated polypeptide 1 (LAP1) deletion led to abnormal VLDL secretion and hepatic steatosis in mice on a chow diet (23). Prior studies showed that hepatocyte-specific deletion of *Lmna* (encoding lamin A/C) caused male-specific hepatic steatosis in C57BL/6J mice, which progressed to NASH and fibrosis after high-fat diet (HFD) feeding (24). This was associated with aberrant expression of genes regulating lipid metabolism, inflammation, and fibrosis in *Lmna*-KO mice compared to controls.

Similarly, a study of a small cohort of twins and siblings with NAFLD identified multiple variants in the gene encoding lamina-associated polypeptide-2α (LAP2α) (25), which is known to bind to lamin A/C and regulate its solubility and distribution within the nucleus, between the lamina and the nucleoplasm (26, 27). In mice harboring global *Lap2α* deletion, cardiac and skeletal muscle abnormalities were noted, but liver development and early (4 week) liver histology were normal (28). However, the role of LAP2α in the protection from, or susceptibility to, NAFLD has not been tested directly.

Thus, despite strong evidence implicating nuclear lamins and their associated proteins in lipid metabolism and human disease, the *in vivo* function and physiologic relevance of LAP2α in hepatocytes remain unknown. To directly test the effect of LAP2α on susceptibility to NAFLD, we have generated mice harboring hepatocyte specific *Lap2α* deletion. Here we report that, unlike *Lmna* and *Lap1* deletion, *Lap2α* deletion protects against HFD-induced steatosis in mice, with reversal of the transcriptional signature seen in *Lmna-KO* mice, including genes associated with lipid metabolism, inflammation, and fibrosis. These data suggest that targeting of lamin-associated proteins, particularly LAP2α, might offer a potential therapeutic strategy in human NASH.

## Results

### Normal baseline histology, with no predisposition to NAFLD, in mice with hepatocytespecific *Lap2a* deletion

Hepatocyte-specific deletion of either *Lmna* or *Lap1* led to spontaneous NAFLD in mice (23, 24), which was male-selective in the former and in both sexes in the latter. We previously reported variants in *TMPO*, encoding LAP2, in twins and siblings with NAFLD (25). Given that the α-isoform of LAP2 is known to regulate lamin A/C, we hypothesized that loss of LAP2α might affect susceptibility to NAFLD in mice. To test this hypothesis, we generated mice with hepatocyte-specific deletion of *Lap2α* (*Lap2α*^(ΔHep)^) as described in Methods. Real-time qPCR (RT-qPCR) of whole liver RNA confirmed reduction in *Lap2α* transcript levels to <5% of WT level in *Lap2α*^(ΔHep)^ mice (**Figure 1A**). *Lap2a*^(ΔHep)^ mice fed chow diet exhibited normal body mass as well as liver morphology and histology, consistent with a prior report (**Figure 1B and C**). Serum ALT and TG levels were normal in both *Lap2α*^(ΔHep)^ and WT mice (**Figure 1D**). These results are in stark contrast to hepatocyte-specific *Lmna* and *Lap1* deletion, which both led to spontaneous hepatic steatosis on chow diet (23, 24).

**Figure 1.**
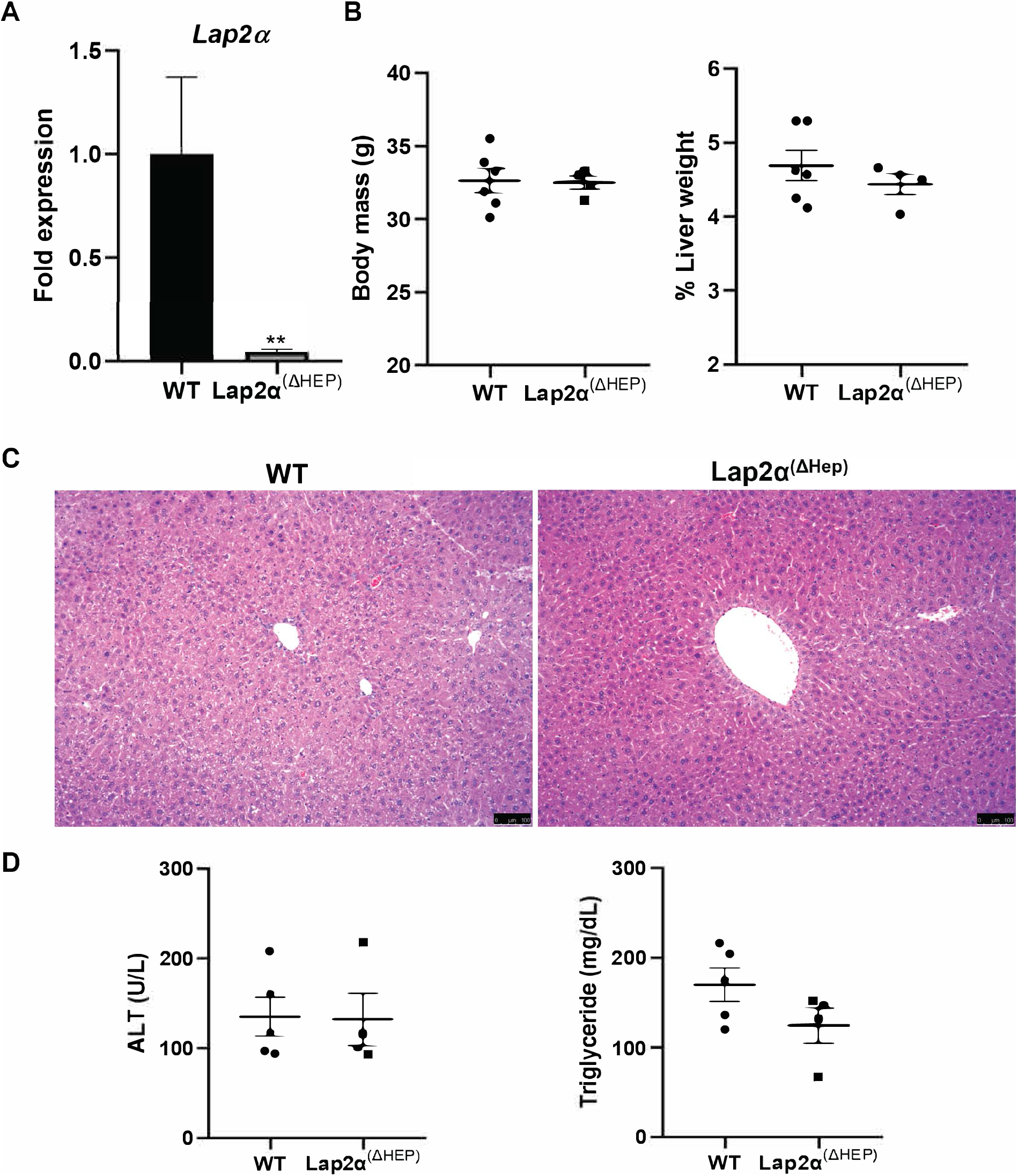
Normal liver histology and serum chemistries in *Lap2α*^(ΔHep)^ mice. (A) *Lap2α* transcript levels were determined via qRT-PCR from whole liver RNA from WT and *Lap2α*^(ΔHep)^ mice, with *Lap2α* transcript detected at <5% of WT levels in *Lap2䎱*^(ΔHep)^ mouse livers (*n*=5 mice per group). (B) Body weight and percentage liver weight of male WT and *Lap2α*^(ΔHep)^ mice. (C) H&E staining of representative liver sections (100X) of male WT and *Lap2α*^(ΔHep)^ mice fed with chow diet (*n*=4-5 mice per group); scale bar, 100 μm. (D) ALT and TG levels in serum. Data represented as mean ± S.E.M. \**P* < 0.05, \*\**P* < 0.01 or \*\*\**P* < 0.001, *Lap2α*^(ΔHep)^ versus WT.

### Loss of *Lap2α* protected against diet-induced hepatic steatosis

To systematically test the role of LAP2α in HFD-induced NAFLD, we subjected *Lap2α*^(ΔHep)^ and control mice to high-fat diet with supplemental sucrose (HFD) for short-term (8 weeks) and long-term (6 months) treatment conditions. As the phenotype in mice with hepatocyte-specific *Lmna* deletion (*Lmna*-KO mice) was most prominent in male mice and we observed only modest hepatic steatosis in female WT and *Lap2α*^(ΔHep)^ mice after 6 months of HFD (**Supplementary Figure 1**), male mice were used for all further testing. After eight weeks of HFD, no increased susceptibility of male *Lap2α*^(ΔHep)^ mice to NAFLD or NASH was observed. Rather, we observed decreased lipid deposition in *Lap2α*^(ΔHep)^ livers compared to WT as determined by oil red O (ORO) staining (**Figure 2A**); blinded semi-quantitative steatosis scoring of hematoxylin and eosin stained sections from the same mice by an expert pathologist showed a trend toward less steatosis in *Lap2α*^(ΔHep)^ mice, though this was not statistically significant. However, after 6 months of HFD, *Lap2α*^(ΔHep)^ livers were found to have significantly decreased hepatic steatosis as determined by ORO staining or by blinded semi-quantitative steatosis scoring (**Figure 2B**).

**Figure 2.**
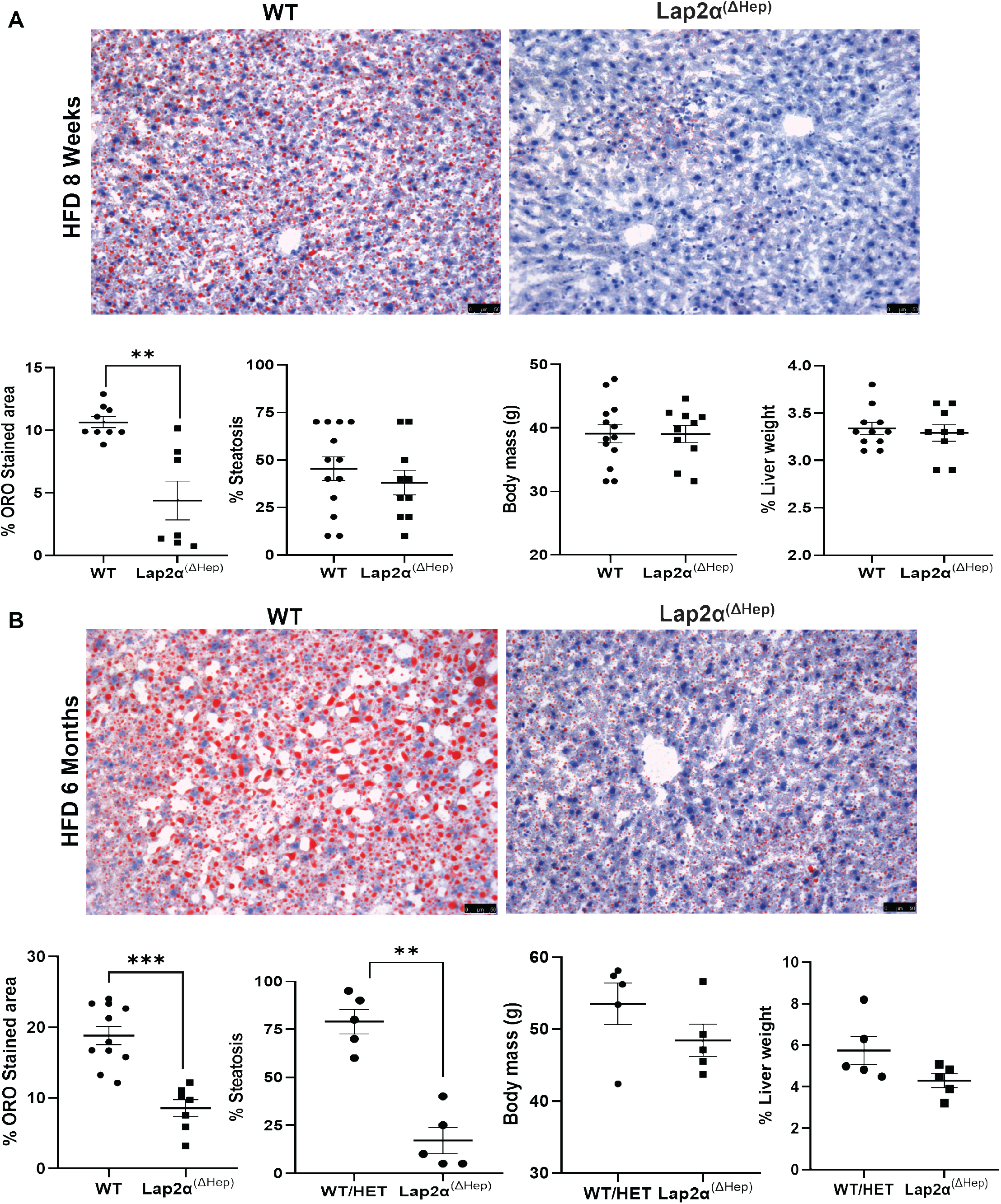
*Lap2α*^(ΔHep)^ mice are protected against HFD diet-induced hepatic steatosis. (A) ORO staining of WT and *Lap2α*^(ΔHep)^ livers after 8 weeks of HFD. Percent steatosis was scored by an expert pathologist in blinded fashion. (B) ORO staining of WT and *Lap2α*^(ΔHep)^ livers after 6 months of HFD, with percent steatosis scored by an expert pathologist in blinded fashion. For (A) and (B), quantitation of ORO was performed as described in Methods, and representative images from two stained livers per genotype per condition are shown. Data represented as mean ± S.E.M. \**P* < 0.05, \*\**P* < 0.01 or \*\*\**P* < 0.001, *Lαp2α*^(ΔHep)^ versus WT; scale bar, 50 μm.

### Loss of *Lap2α* did not affect lamin A/C expression

Given that *Lmna* deletion resulted in NAFLD (24) and that loss of *Lap2α* appeared to protect against NAFLD (Figure 2), and the known role of LAP2α in the regulation of lamin A/C, we asked whether lamin A/C expression or distribution within the nucleus might be altered in *Lap2α*^(ΔHep)^ livers. It was previously reported that nucleoplasmic LAP2α binds to lamin A/C, regulating its mobility and assembly state (27, 29). To examine the distribution of lamin A/C in the absence of hepatocyte LAP2α, we performed immunofluorescence staining of mouse liver tissue. Lamin A/C staining at the nuclear rim was overall not different in *Lap2α*^(ΔHep)^ livers compared to WT, though we observed particularly prominent lamin A/C staining in *Lap2α*^(ΔHep)^ mice after 8 weeks of HFD (**Figure 3A**); under chow diet conditions, lamin A/C staining was also similar in WT and *Lap2α*^(ΔHep)^ mouse livers (**Supplementary Figure 2**). Consistent with prior reports (27, 29), immunoblot analysis of whole livers did not show significantly different overall lamin A/C levels in livers from *Lap2α*^(ΔHep)^ and WT mice (**Figure 3B, Supplementary Figure 2**).

**Figure 3.**
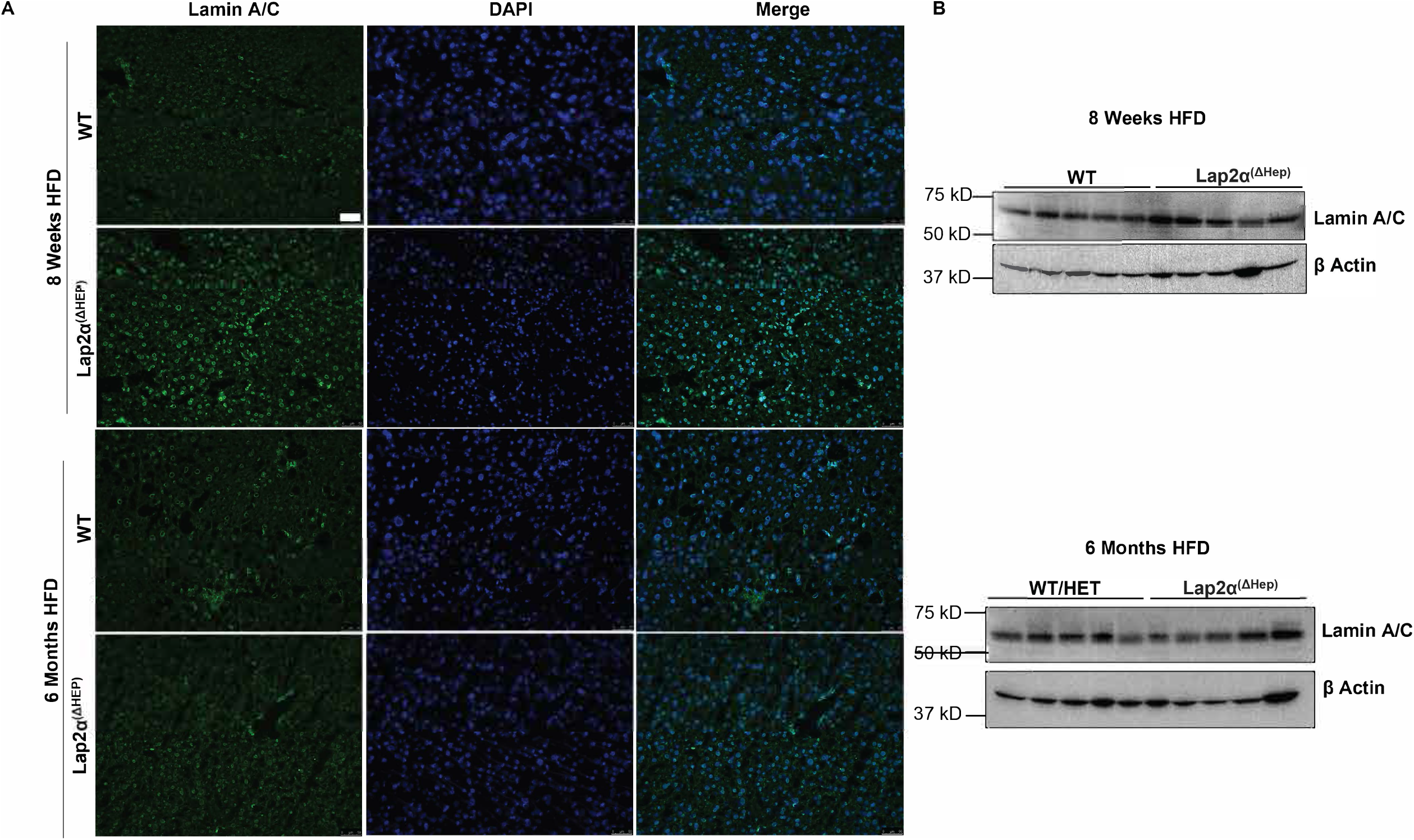
Unchanged lamin A/C expression and nuclear rim staining in *Lap2α*^(ΔHep)^ mouse livers. (A) Livers of 3 WT and 3 *Lap2α*^(ΔHep)^ mice after short term (8 weeks) and long term (6 months) of HFD were cryosectioned and stained for lamin A/C and DAPI. Images were acquired at 100X with a Leica DMRB 5000B microscope, and representative images are shown; scale bar, 50 μm. (B) Protein from livers of WT and *Lap2α*^(ΔHep)^ mice fed short and long-term HFD was extracted and analyzed by immunoblotting using anti-lamin A/C and β-actin antibodies (note that the anti-lamin antibody preferentially recognizes mouse lamin C compared to lamin A). Each lane corresponds to an individual liver.

### Pro-steatotic genes were downregulated in *Lap2α*^(ΔHep)^ mice

Given that *Lap2α*^(ΔHep)^ mice were protected against HFD-induced NAFLD, as well the known role of LAP2α in regulating lamin A/C distribution and association with chromatin (27), we hypothesized that the pro-steatotic transcriptional changes seen in *Lmna*-KO mice might be reversed in the setting of *Lap2α* deletion. To evaluate this hypothesis, we selected some of the most highly up-regulated pro-steatotic genes in *Lmna-KO* mice, including *Cidea, Cd36*, and *Mogat1*, for analysis. *Cidea* encodes a member of the CIDE family of proteins, which regulate lipogenesis and lipolysis (30, 31). In contrast to *Lmna*-KO mice, *Lap2α*^(ΔHep)^ mice showed significantly reduced levels of *Cidea* transcript compared to WT mice after 8 weeks and 6 months of HFD (**Figure 4A**, **4B**). Notably, *Cidea* was also highly downregulated in *Lap2α*^(ΔHep)^ mice compared to WT under chow diet conditions (**Figure 4C**), suggesting that this transcriptional difference was a direct result of loss of LAP2α and a contributor to protection from NAFLD, rather than a consequence of decreased steatosis in *Lap2α*^(ΔHep)^ mice. Consistent with these results, levels of *Cd36*, which encodes a fatty acid translocase (32, 33), were decreased in *Lap2α*^(ΔHep)^ mice at baseline and after long-term HFD. Similarly, expression of *Mogat1*, encoding the enzyme responsible for conversion of monoacylglycerol to diacylglycerol (34, 35), was also decreased in *Lap2α*^(ΔHep)^ mice after long-term HFD and at baseline (**Figure 4B**, **4C**). Taken together, these data indicate that loss of LAP2α in hepatocytes protects mice against HFD-induced steatosis, with transcriptional downregulation of pro-steatotic genes.

**Figure 4.**
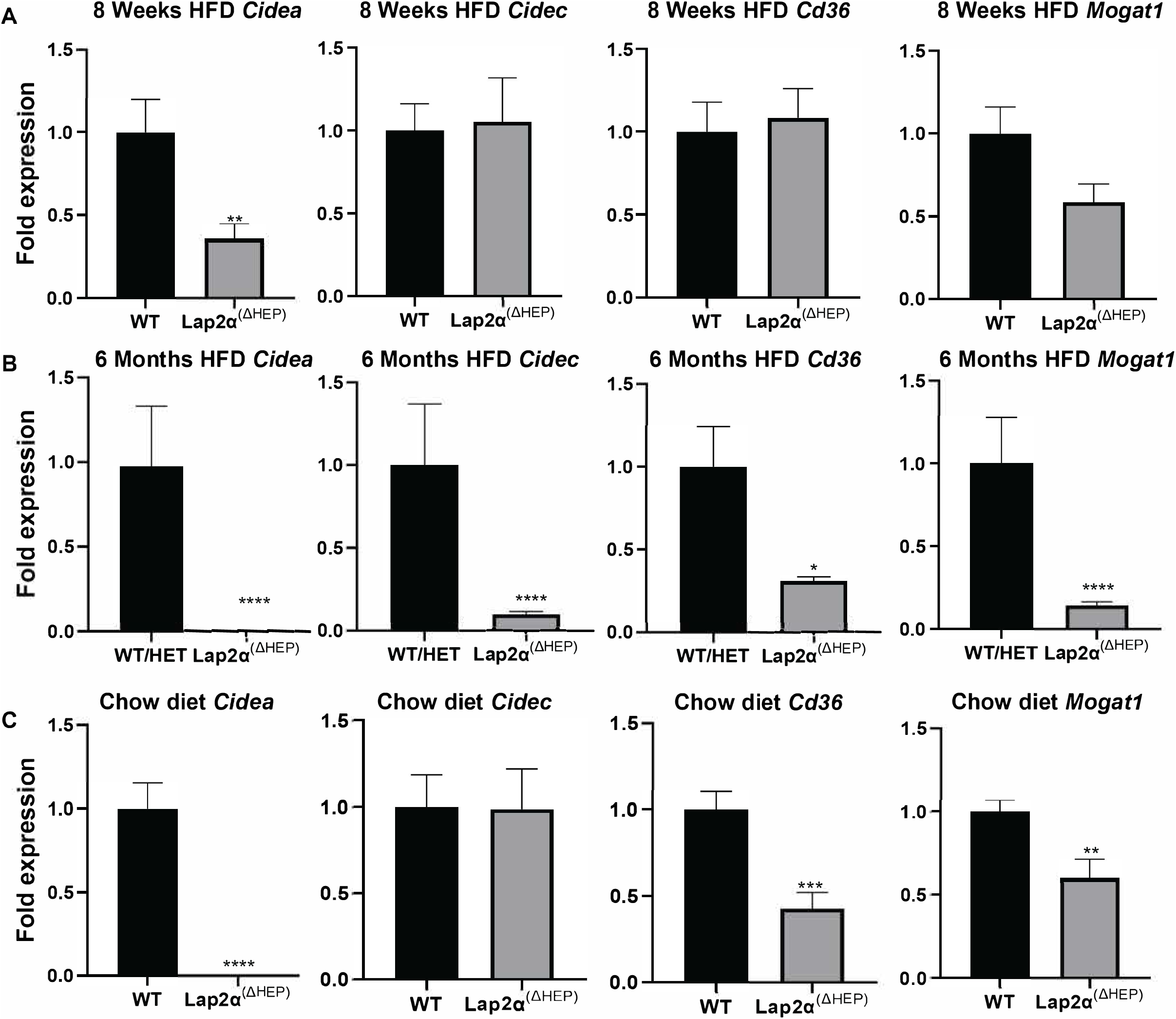
Downregulation of pro-steatotic genes in *Lap2α*^(ΔHep)^ livers. Transcript levels of selected lipid metabolism genes were analyzed via qPCR using RNA from control and *Lap2α*^(ΔHep)^ livers from (A) short-term HFD (8 weeks; *n*=5 mice per group); (B) long-term HFD (6 months; *n*=5 mice per group); (C) Chow diet, *n*=4-6 mice per group. Data represented as mean ± S.E.M. \**P* < 0.05, \*\**P* < 0.01 or \*\*\**P* < 0.001, *Lap2α*^(ΔHep)^ versus control; where indicated, heterozygous mice were included as control mice with WT group.

### Loss of *Lap2α* protected against NASH and decreased expression of pro-inflammatory and pro-fibrotic genes in long-term HFD-fed mice

In human NAFLD/NASH, hepatocyte injury, steatohepatitis, and progressive fibrosis are thought to be the primary mediators of long-term sequelae including cirrhosis and hepatocellular carcinoma (3, 5). Notably, *Lmna-KO* mice were more susceptible to both inflammation and fibrosis compared to control mice (24). Given that *Lap2α*^(ΔHep)^ mice were protected against hepatic steatosis and that the most highly upregulated pro-steatotic genes in *Lmna*-KO mice were downregulated in the absence of LAP2α, we hypothesized that *Lap2α*^(ΔHep)^ mice might be protected against NASH, inflammation, and early fibrosis compared to control mice. Among all mice fed chow diet or short-term (8 weeks) HFD, no fully developed NASH was seen in either *Lap2cα*^(ΔHep)^ or WT mice (**Figure 5A**). However, among mice fed HFD for 6 months, although serum ALT and TG levels did not differ, NAFLD activity scores (NAS, as previously described (36), with slight modification as per Methods section) were significantly lower in *Lap2α*^(ΔHep)^mice compared to controls (**Figure 5B**), suggesting that loss of LAP2α protected against lipid-mediated hepatocyte injury and inflammation in addition to hepatic fat deposition. Consistent with this, *Lap2α*^(ΔHep)^ mice showed a trend toward reduced expression of pro-inflammatory genes including *Ubd, Irf7, Stat1, Themis*, and *Tnfa*, though for some of these genes the difference did not reach statistical significance (**Figure 6A**); all of these genes had been highly upregulated in *Lmna*-KO mice (24). As expected with HFD lacking high cholesterol and fructose content (37, 38), we did not observe any significant fibrosis in any mice under any dietary condition. Reassuringly, in livers from *Lap2α*^(ΔHep)^ mice after 6 months of HFD, as compared to control livers, there was no increase in expression of pro-fibrotic genes; rather, there was a trend toward decreased expression of several pro-fibrotic genes, including *Tgfb*, *Col1a1*, *Timp1*, and *Acta2*, though this did not reach statistical significance (**Figure 6B**).

**Figure 5.**
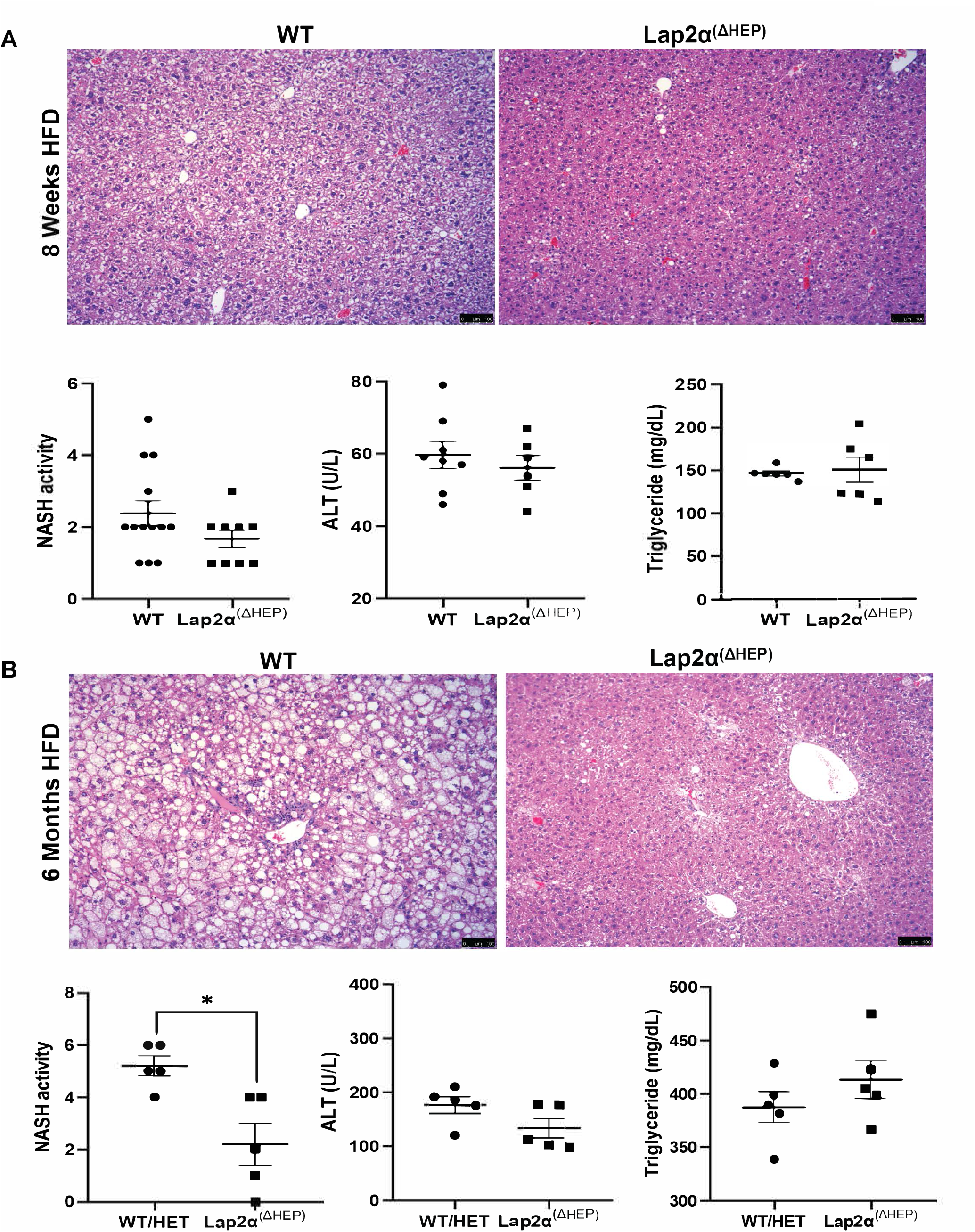
Decreased steatohepatitis in *Lap2α*^(ΔHep)^ livers. H&E staining of liver sections of control and *Lap2α^(ΔHep)^* mice was performed, and images were acquired via Leica DMRB 5000B microscope, with representative images shown. NAFLD activity scores (NAS) were determined by an expert liver pathologist in blinded fashion, and serum ALT and TG were measured. (A) Mice were fed with short term HFD (8 weeks; *n*>9 mice per group); (B) Mice were fed with long-term HFD (6 months; *n*=5 mice per group). Data represented as mean ± S.E.M. \**P* < 0.05, \*\**P* < 0.01 or \*\*\**P* < 0.001, *Lap2α^(AHep)^* versus control; where indicated, heterozygous mice were included as control mice with WT group. Scale bar, 100 μm.

**Figure 6.**
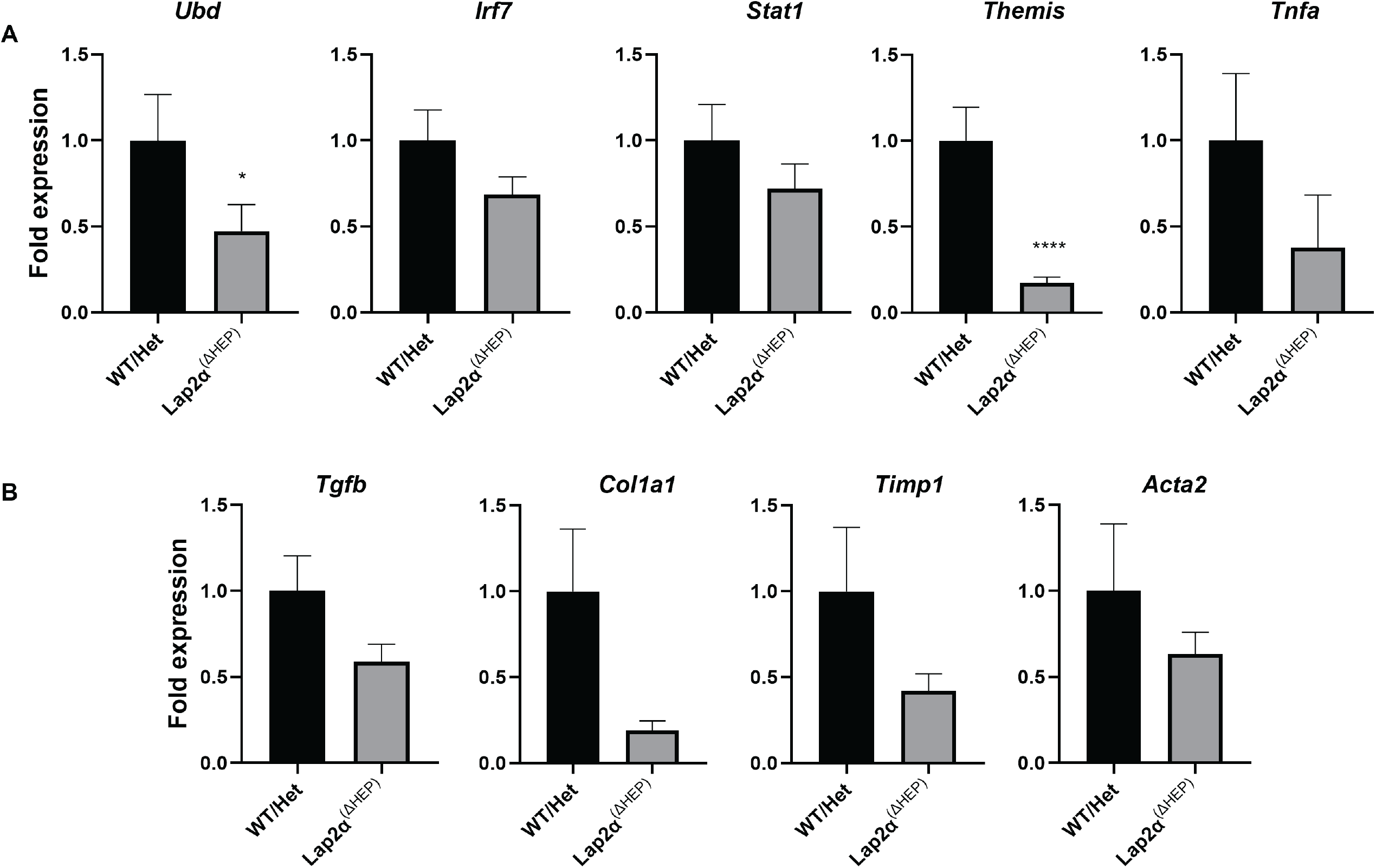
Downregulation of select pro-inflammatory and no increase in pro-fibrotic genes in *Lαp2α*^(ΔHep)^ livers. Transcript levels of selected pro-inflammatory genes (A) and pro-fibrotic genes (B) that were highly upregulated in *Lmna-KO* mice (24) were determined using whole liver RNA from control and *Lap2α*^(ΔHep)^ mice fed HFD for 6 months (*n*=5 mice per group). Data represented as mean ± S.E.M. \**P* < 0.05, \*\**P* < 0.01 or \*\*\**P* < 0.001, *Lap2α*^(ΔHep)^ versus control; as indicated, heterozygous mice were included as control mice with WT group.

## Discussion

Mutations in genes encoding nuclear envelope proteins are known to cause lipodystrophy syndromes with hepatic steatosis and progression to NASH (13, 21, 39–41). Evidence is now accumulating from genetic animal models for the functional importance of the hepatocyte nuclear envelope in NAFLD (23, 24, 42). Hepatocyte-specific deletion of either *Lmna* or *Lap1* leads to hepatic steatosis in mice, though the phenotypes of the two models appear to be distinct, with a male-predominant phenotype in the case of *Lmna* deficiency and a prominence of nuclear lipid droplets in the case of *Lap1* deficiency.

Herein we show that, in contrast to *Lmna* or *Lap1* deletion, hepatocyte-specific *Lap2α* deletion protected male mice from high fat diet-induced NAFLD and NASH. It was previously shown that LAP2α binds to lamin A/C via the proteins’ respective carboxy-terminal tails, with consequent regulation and maintenance of lamin A/C in the nuclear interior in a mobile and low assembly state (26, 27). These protein-protein interactions influence several physiological functions, including proliferation and differentiation, as well as chromatin organization and gene expression (43–45). Our data, in agreement with prior reports (27, 29), show that loss of LAP2α had no effect on lamin A/C expression overall, with a variable effect on lamin A/C staining at the nuclear periphery. Nevertheless, there were transcriptional changes in pro-steatotic, pro-inflammatory, and pro-fibrotic genes that were opposite to those seen in *Lmna*-KO mice. In particular, whereas male mice lacking lamin A/C in hepatocytes were predisposed to steatohepatitis via a dramatic increase in transcriptional expression of pro-steatotic genes including *Cidea, Cidec, Mogat1*, and *Cd36* (24), in this study we observed opposite changes in the expression of all of these genes in the absence of LAP2α. While lamin A/C-independent mechanism(s) of protection via loss of LAP2α cannot be ruled out, these data suggest the possibility that the observed protection in LAP2α-deficient mice could be due to enhanced hepatoprotective lamin A/C-mediated regulation of gene expression in hepatocytes. Our results are in alignment with previous findings and support the idea that lamin A/C has different properties at the nuclear periphery versus within the nucleoplasm and that it is regulated by LAP2α (27, 29, 46). Additionally, our data are supportive of a model in which there are mechanistic differences between LAP1-related steatosis (male equal to female, prominent nuclear lipid droplets) and lamin A/C and LAP2α-related steatosis (male > female, nuclear lipid droplets not prominent). This may reflect a predominance of ER dysfunction and defective lipid secretion in the case of loss of LAP1 (23, 42) versus a predominantly gene regulation-based mechanism of susceptibility to NAFLD in the case of lamin A/C and LAP2α (24).

It is important to note that variants in *TMPO*, encoding the six LAP2 isoforms including LAP2α, associated with increased risk of NAFLD in a twin and sibling cohort (25), rather than protection from NAFLD. However, some of these variants were predicted to impact multiple LAP2 isoforms, rather than the α-isoform specifically, and thus their potential impacts cannot be directly compared to the effects of *Lap2α* deletion in the current study. Additionally, the consequences of *TMPO* variants that impact LAP2α function outside of its interaction with lamin A/C may be difficult to predict, while variants that enhance the LAP2α-lamin A/C interaction would be predicted to increase the risk of NAFLD. Finally, such germline genetic variants could impact both liver development and extrahepatic metabolism in ways that hepatocyte-specific deletion of the α-isoform of LAP2 via an albumin-*Cre* transgene (47), as in the current study, did not. Together, these differences may account for the apparently contradictory effects of *TMPO* variants in humans and *Lap2α* deletion in mice.

Collectively, our findings advance the current understanding of the physiological roles of LAP2α, lamin A/C, and the nuclear envelope in the onset and progression of steatosis and NASH. Notably, whereas *Lmna-KO* mice were more susceptible to NASH and fibrosis compared to controls (24), here we observed significant protection from steatohepatitis in *Lap2α*-deficient (*Lap2α*^(ΔHep)^) mice compared to WT and heterozygous control mice, with decreased expression of pro-inflammatory genes and no increase (with a potential trend toward decrease) in pro-fibrotic genes. This apparent protection provided by loss of LAP2α from not only hepatic steatosis, but also NASH and susceptibility to fibrosis, is important given that morbidity and mortality of NAFLD are highly correlated with steatohepatitis and fibrosis (3, 5). Taken together, our data suggest that hepatocyte LAP2α may be a potential therapeutic target to enhance repression of lipogenic, pro-inflammatory, and pro-fibrotic gene expression in humans to prevent the development and/or progression of NASH.

## Methods

### Antibodies

For immunoblot, anti-lamin A/C (1:1000, catalog number 2032, Cell Signaling Technology, Beverly, MA), anti-β-actin (1:1000, Cell Signaling Technology 4970), and anti-rabbit IgG-HRP secondary antibody (1:1500, Sigma Aldrich, St. Louis, MO) were used. For immunofluorescence, anti-lamin A/C (1:100, sc-376248, Santa Cruz Biotechnology, Santa Cruz, CA) and Alexa Fluor 488 rabbit anti-mouse IgG (H+ L) (1:200, A11059, Thermo Scientific) were used.

### Animal experiments

C57BL/6J mice with *Lap2α* specific exon 4 flanked by loxP sites on chromosome 10 were described previously (29); these mice were crossed with C57BL/6J mice expressing a transgene with Cre recombinase expression governed by the albumin promoter (Jackson Laboratories (47)) to generate C57BL/6J offspring with hepatocyte specific deletion of exon 4 of *Lap2α* and littermate control mice lacking the albumin-*Cre* transgene (or where indicated, lacking one floxed *Lap2α* allele – such mice were considered to be heterozygous). Male *Lap2α*^(ΔHep)^ (Cre^+^, *Lap2α ^flox/flox^*), wild-type (WT) control (Cre^-^, *Lap2α ^flox/flox^*) mice, and where indicated, heterozygous (Cre^+^, *Lap2α ^flox/WT^*) mice (8-10 weeks of age) were used for further studies. Mice were fed normal chow or HFD (58% fat calories, 18% sucrose by weight, 23% protein by weight; Research Diets D12331, New Brunswick, NJ) for 8 weeks or up to 6 months as indicated.

For the 8-week HFD challenge, whole blood was collected by intracardiac puncture, and the liver was harvested under isoflurane anesthesia. For normal chow diet and 6-month HFD, mice were euthanized by CO_2_ asphyxiation prior to cardiac puncture and harvesting of the liver. Whole blood was centrifuged at 4°C and 3000 RPM for 10 min for serum collection. Liver tissue was stored in 10% formalin (for histology), optimum cutting temperature compound (OCT) for immunofluorescence staining, RNAlater (for gene expression studies) or snap-frozen and stored at −80°C (for protein analysis). Mice and whole livers were weighed before liver processing to calculate the liver percentage of body mass (% liver weight).

### Biochemical Parameters and Liver histology

Serum alanine aminotransferase (ALT) and serum triglyceride (TG) values were determined by the Unit for Laboratory Animal Medicine at the University of Michigan. Paraffin-embedded livers were cut into 6 μm sections and stained with hematoxylin and eosin (H&E); images were captured using a Leica DM 5000B microscope. An expert liver pathologist (E.K.C.) scored the stained sections in a blinded fashion for steatosis and NASH activity, with the latter according to the method of Kleiner et al. (36), with slight modification. Briefly, microvesicular steatosis was included together with macrovesicular steatosis in the histologic scoring, and slightly lower thresholds were used to assign lobular hepatitis scores compared to the original description.

### Oil Red O Staining (ORO) and Quantitation

Six-micron thick frozen liver sections were fixed in ice-cold 10% formalin and washed three times with water, followed by 5 mins in absolute propylene glycol (Sigma Aldrich) and stained with 0.5% ORO (Sigma Aldrich) for 10 min at 60°C. Stained slides were washed and counterstained for 30-45 seconds with Gill’s 3 Hematoxylin (Sigma Aldrich), then rinsed thoroughly with water and mounted with glycerol gelatin (Sigma Aldrich). Eight to ten fields per liver section (10X objective) were photographed using a Leica DM 5000B microscope for analysis. Fiji ImageJ Analysis Software version 1.51j8 (National Institutes of Health, Bethesda, MD) was used to calculate the proportion of each section with positive ORO staining.

### Immunofluorescence Staining

OCT-embedded frozen liver tissue was cut into 6-micron sections and fixed in methanol at −20 °C for 10 min, followed by washed, permeabilization (0.1% Triton X-100 in PBS), and blocking (5% bovine serum albumin in PBS). Primary antibody was incubated overnight at 4 °C followed by a 1-hour incubation with Alexa Fluor–488 goat anti-mouse IgG. Slides were mounted using Prolong Gold Anti-Fade Reagent with DAPI, and stained sections were visualized with a Leica DM 5000B fluorescence microscope.

### Quantitative real-time polymerase chain reaction (qPCR) analysis

Total RNA from WT and *Lap2α*^(ΔHep)^ mice was isolated using RNeasy Mini Kit (Qiagen), and cDNA was synthesized using iScript cDNA Synthesis kit (BIO-RAD CA, USA). Transcript levels of genes of interest were quantified using StepOne Real time PCR system (Thermo Scientific, CA, USA); qPCR primer sequences are shown in **Supplementary Table 1**. Relative expression was determined after normalizing to 18S RNA and use of 2^-ΔΔCT^ method.

### Immunoblotting

Liver samples were homogenized using T-PER tissue protein extraction reagent buffer (Thermo Scientific, CA) containing protease and phosphatase inhibitors (Sigma). Total protein was quantified using BCA Kit (Thermo Scientific), and equal amount of protein was separated on 4-12% Novex tris-glycine gels (Thermo Scientific). Proteins were transferred to PVDF membrane (Bio-Rad, USA) and analyzed with anti-lamin A/C antibody. Blots were stripped using stripping buffer (Thermo Scientific) and probed with β-actin antibody (1:1500) to confirm equivalent protein loading.

### Statistics

Data are expressed as mean ± SEM and analyzed by unpaired t test (2-tailed) or one-way analysis of variance (ANOVA), followed by the Mann-Whitney test using Graph Pad Prism 9.0 (CA, USA). \**P*<0.05, \*\**P*<0.01 and \*\*\**P*<0.001 were considered to be significant.

## Study approval

Mouse experiments were performed in accordance with guidelines outlined in the Guide for the Care and Use of Laboratory Animals prepared by the National Academy of Sciences and published by the National Institutes of Health, and with approval from the University of Michigan Institutional Animal Care and Use Committee (Protocol numbers PRO00009549 (G.F.B.) and PRO00010138 (Mouse Metabolic Phenotyping Center)).

## Author Contributions

K.K.U., M.B.O., and G.F.B. conceived and designed the study; K.K.U. and G.F.B. performed the experiments; E.K.C. performed the histologic scoring of mouse liver sections; R.F. provided the floxed *Lap2α* mice; K.K.U. and G.F.B wrote the manuscript; all authors critically reviewed and revised the manuscript.

## Acknowledgment

The work was supported by National Institutes of Health (NIH) grants R01 DK116548 (M.B.O.), K08 DK120948 (G.F.B.), and U2CDK110768 (University of Michigan Mouse Metabolic Phenotyping Center), and by the University of Michigan Department of Pediatrics Equipment Core.

