## Supplementary Material for "Hepatocyte-specific loss of LAP2α protects against diet-induced hepatic steatosis and steatohepatitis in male mice"

### Supplementary Figure 1

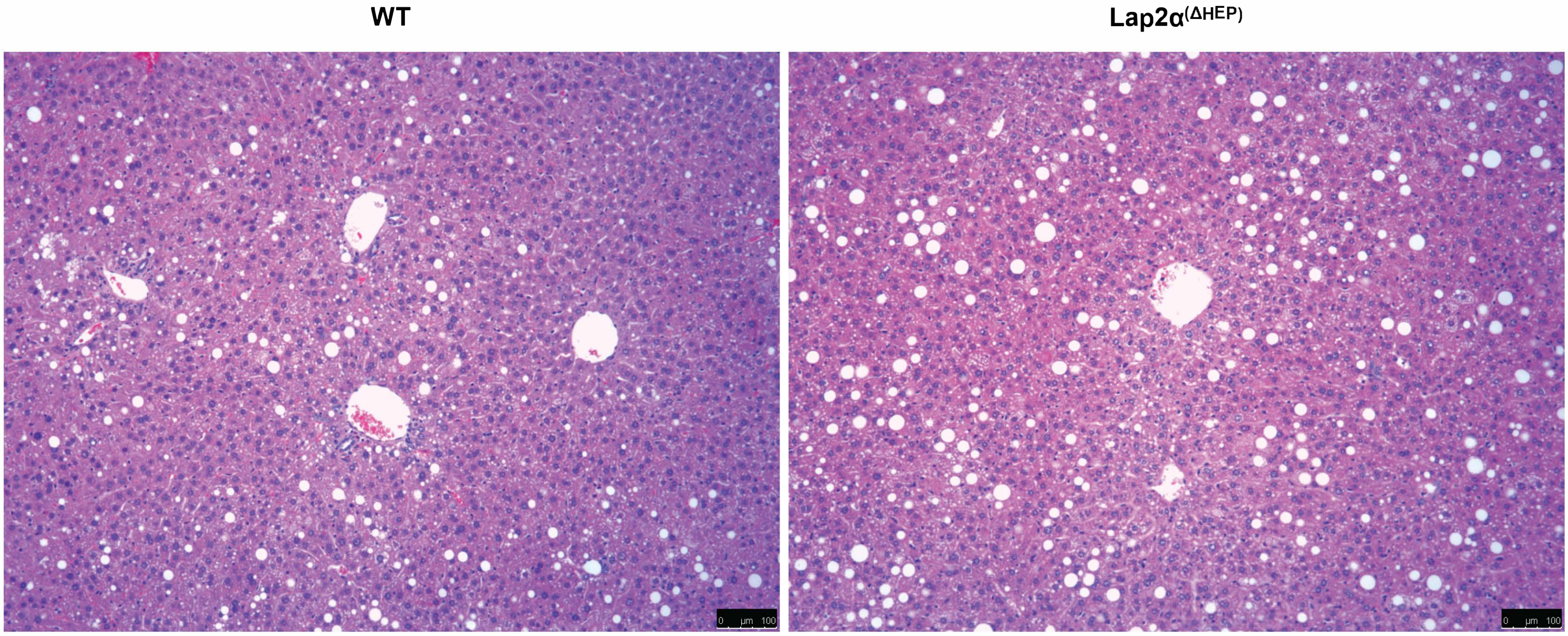


**Supplementary Figure 1. Modest steatosis in female WT and *Lap2α*$^{(\Delta Hep)}$ mice after 6 months of HFD.** H&E staining of representative liver sections (100X) of female WT and *Lap2α*$^{(\Delta Hep)}$ mice fed with HFD for 6 months (*n*=3-4 mice per group) showing modest hepatic steatosis; scale bar, 100 µm.

### Supplementary Figure 2

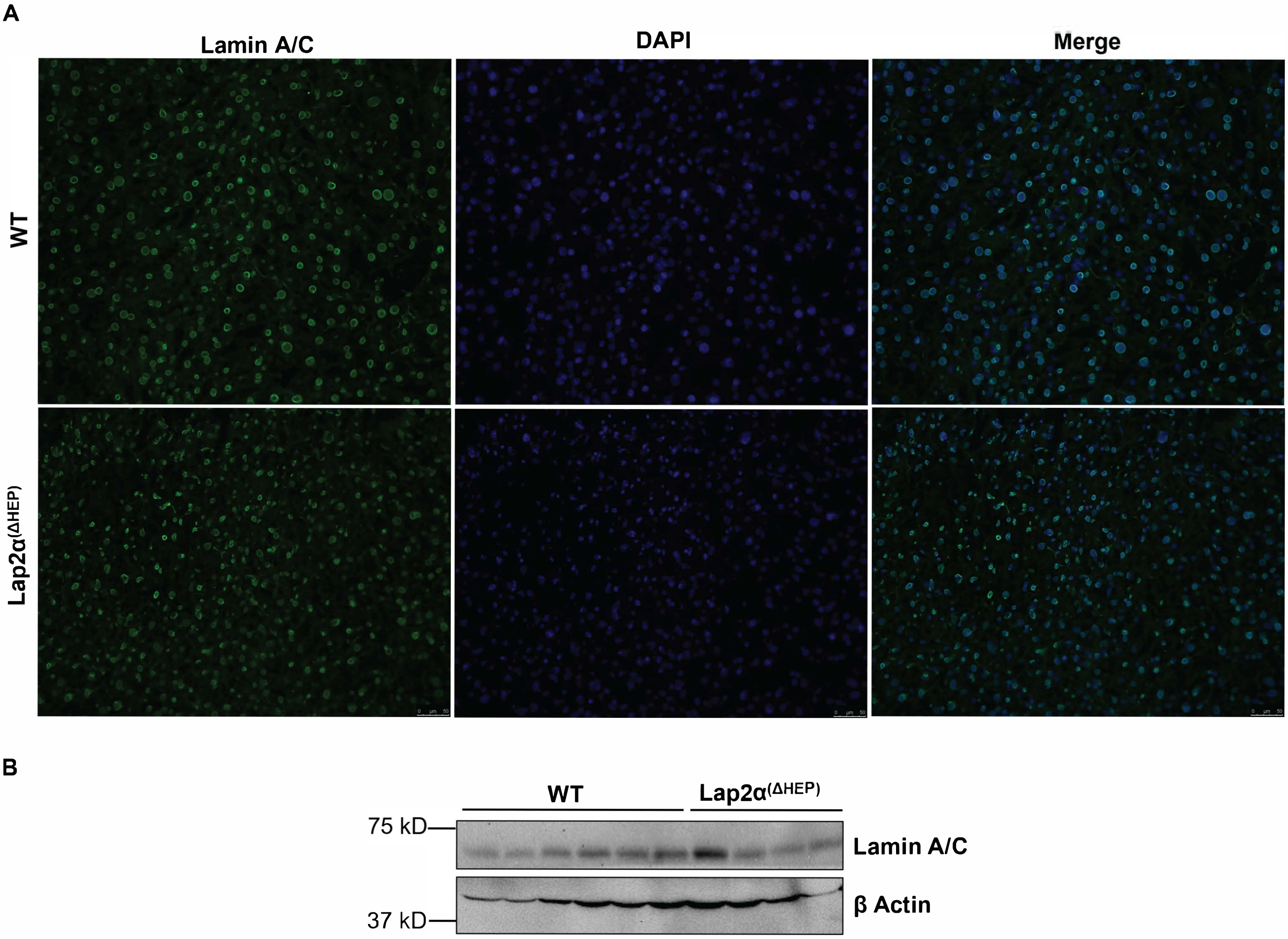


**Supplementary Figure 2. Lamin A/C staining in liver sections from WT and *Lap2α*$^{(\Delta Hep)}$ mice fed chow diet.** (A) Livers from 3 WT and 3 *Lap2α*$^{(\Delta Hep)}$ male mice after chow diet were cryosectioned and stained for lamin A/C and DAPI. Images were acquired at 100X with a Leica DMRB 5000B microscope, and representative images are shown; scale bar, 50 µm. (B) Whole lysates of livers of WT and *Lap2α*$^{(\Delta Hep)}$ mice fed chow diet were analyzed by immunoblotting using anti-lamin A/C and β-actin antibodies. Each lane corresponds to an individual liver.

| Gene | Forward (5′→3′) | Reverse (5′→3′) |
|---|---|---|
| *Lap2α* | TCCTCTACTCCTCTTCCAACAGTC | GCGGACTTCACTTTCTTGTGT |
| *Cidea* | CAGTCTGCAAGCAACCAAAG | TTGTGCATCGGATGTCGTAG |
| *Cidec* | AGCTAGCCCTTTCCCAGAAG | CCTTGTAGCAGTGCAGGTCA |
| *Cd36* | GCTGCACCACATATCTACCAAA | AGGATATGGAACCAAACTGAGG |
| *Mogat1* | TCAAAACGCAGGATTTGGAT | ACAACGGGAAACAGAACCAG |
| *Ubd* | CCTTACCCTGAAGGTGGTGA | CTTCCAGCTTCTTTCCGTTG |
| *Irf7* | TGATCTTTCCCAGTCCTGCT | TGCCTACCTCCCAGTACACC |
| *Stat1* | GTGGAGCCCTACACGAAAAA | ATACTTCCCAAAGGCGTGGT |
| *Themis* | TGTGAAGGTGGCTGTGAGAG | CATCTGCAGGCAAACTGCTA |
| *Tnfa* | CGTCAGCCGATTTGCTATCT | CGGACTCCGCAAAGTCTAAG |
| *Tgfb* | GGCCAGATCCTGTCCAAACT | TGTTGCGGTCCACCATTAG |
| *Col1a1* | TCTGACTGGAAGAGCGGAGAG | GGCACAGACGGCTGAGTAGG |
| *Timp1* | TCCCTTGCAAACTGGAGAGT | AGGTGCACAAGCCTGGATTC |
| *Acta2* | TCAGCGCCTCCAGTTCCT | AAAAAAAACCACGAGTAACAAATCAA |

**Supplementary Table 1. Primers used in qPCR analysis.** All primer sequences aside from those for *Lap2α* were previously reported (1); *Lap2α* primer sequences were adopted from Stenvall et al. (2).
